# Nuclear genetic regulation of human mitochondrial RNA modification

**DOI:** 10.1101/666339

**Authors:** Aminah T. Ali, Youssef Idaghdour, Alan Hodgkinson

## Abstract

RNA modifications affect the stability and function of RNA species, regulating important downstream processes. Modification levels are often dynamic, varying between tissues and individuals, although it is not always clear what modulates this variation or what impact it has on biological systems. Here, we quantify variation in RNA modification levels at functionally important positions in the mitochondrial genome across 11,552 samples from 39 tissue/cell types and find evidence that modification levels impact mitochondrial transcript processing. We identify novel links between mitochondrial RNA modification levels in whole blood and genetic variants in the nuclear genome, including missense mutations in *LONP1* and *PNPT1*, as well as missense mutations in *MRPP3*, *SLC25A26* and *MTPAP* that associate with RNA modification levels across multiple tissue types. Genetic variants linked to modification levels are associated with multiple disease phenotypes, including blood pressure, breast cancer and Moyamoya disease, suggesting a role for these processes in complex disease.

## Introduction

RNA modifications are post-transcriptional changes to the chemical composition of nucleic acids and represent a means by which RNA function can be fine-tuned (Li and Mason 2014). Sites of RNA modification are often highly evolutionarily conserved and are crucial for processes such as development, cell signalling and maintenance of the circadian rhythm, pointing to a major role for RNA modification in fundamental cellular processes (Li and Mason 2014). To date, over 160 different types of RNA modification have been identified (Boccaletto et al. 2018), occurring on several types of RNA molecule, though they are found most abundantly on ribosomal and transfer RNAs (Machnicka et al. 2013). The exact role of an RNA modification depends on the type, location and target of the modification. Within tRNAs for example, modifications in the anticodon region can increase a tRNAs decoding capacity, and improve translational fidelity (Yarian et al. 2002), whereas modifications to the core of a tRNA molecule can promote correct folding and structural stability (Helm 2006). Modifications to rRNA molecules are largely involved in the stabilisation of the ribosome structure, but can also facilitate protein synthesis (Sloan et al. 2017), and modifications to mRNA molecules can affect the maturation, translation and degradation of an mRNA molecule by either recruiting additional proteins or by altering the secondary structure of the mRNA (Zhao et al. 2017). Importantly, not all modifications are fixed; instead, some display a dynamic range of modification (Hauenschild et al. 2015), which may in turn reflect a dynamic mode of RNA regulation.

Interest in RNA modifications has recently been renewed, due to the development of high-throughput technologies that can detect modifications on a transcriptome-wide scale. However, these studies are often small in scale and limited to specific cell lines, and frequently focus on the detection of novel modification sites rather than attempt to survey the dynamic range of modification level across a large number of sites and samples (Clark et al. 2016; Dominissini et al. 2016; Li et al. 2017; Safra et al. 2017). Here, we utilise computational methods (Sanchez et al. 2011; Hodgkinson et al. 2014; Hauenschild et al. 2015; Idaghdour and Hodgkinson 2017) to quantify RNA modification levels across a total of 11,552 human RNA sequencing (RNAseq) libraries in 39 tissue/cell types. Within this, we focus on data from mitochondrial-encoded RNA, where due to its abundance across all tissues, we can detect RNA modifications at multiple functionally important positions along the mitochondrial transcriptome.

Mitochondria have essential roles in multiple cellular processes, including energy production, signalling, ion metabolism and apoptosis. The mitochondrial genome encodes 2 rRNA genes, 22 tRNA genes, and 13 mRNA genes (Anderson et al. 1981), and is transcribed polycistronically, after which it is processed according to the ‘tRNA punctuation model’ to release individual RNA components (Ojala et al. 1981), whereby tRNAs interspersed between rRNA and mRNA genes are cleaved by nuclear-encoded proteins (Holzmann et al. 2008; Brzezniak et al. 2011; Sanchez et al. 2011; Powell et al. 2015). Intermediate and mature RNA transcripts harbour extensive RNA modifications, which impact features such as transcript structure and stability that can be important for both processing and function (Rorbach and Minczuk 2012). Interestingly, steady state levels of mature mitochondrial transcripts vary substantially from the 1:1 ratio that might be expected from polycistronic transcription (Mercer et al. 2011), indicating the importance of post-transcriptional processes in the maintenance of mitochondrial homeostasis.

In illustration of this, knock-down of nuclear encoded mitochondrial RNA processing enzymes in mice leads to the accumulation of unprocessed mitochondrial encoded transcripts, decreased levels of protein synthesis and altered mitochondrial respiration rates (Sanchez et al. 2011; Sen et al. 2016). Likewise, altered modification of mitochondrial-encoded RNA can have similar consequences; 1) methylation of bases on the ninth position of certain mitochondrial tRNAs is understood to have an impact on tRNA structure and stability (Helm et al. 1999; Helm 2006) which in turn may influence post-transcriptional tRNA recognition and cleavage dynamics and therefore downstream protein translation; 2) methylation of *mt-RNR2* transcripts at mtDNA position 2617 is believed to provide stabilising interactions to mature mitoribosomes, and lack of methylation at this position has been linked to impaired mitochondrial protein synthesis (Bar-Yaacov et al. 2016); 3) methylation of *mt-ND5* transcripts at mtDNA position 13710 varies according to tissue type (Safra et al. 2017), and interferes with translation through mitoribosome stalling and leads to decreased protein levels (Li et al. 2017). In this study, we focus on quantifying variation in mitochondrial encoded RNA modification levels at these three classes of site on a population level across multiple tissue types. We perform quantitative trait mapping using mitochondrial-encoded RNA modification rates in order to identify nuclear genetic variants and genes that are involved in the regulation of these processes across tissues, and to unravel their downstream functional consequences.

## Results

### Overview

In order to study variation in modification levels of mitochondrial-encoded RNA across multiple tissue types, we mapped and filtered 13,857 RNAseq samples from 39 tissue/cell types, across 5 independent datasets (see Methods and Materials, Supplementary Table 1), to the human reference genome using a stringent pipeline optimised for the analysis of mitochondrial data. Previous work has demonstrated that the levels of a particular form of mitochondrial RNA modification (RNA methylation) can be inferred at particular nucleotides using the proportion of mismatching bases in RNAseq data (Mercer et al. 2011; Sanchez et al. 2011; Hauenschild et al. 2015) (see Methods and Materials for details). As such, we use this proportion to quantify the level of RNA methylation at three categories of modified site where RNA methylation is known to be functionally important; 1a) methylation at position 9 (henceforth referred to as P9 sites) of 13 different mt-tRNAs along the mitochondrial genome, 1b) an averaged estimate of methylation level across 11 different mt-tRNA P9 sites that consistently show variation in whole blood (Hodgkinson et al. 2014), 2) methylation at mt-DNA positon 2617 within *mt-RNR2* and 3) methylation at mt-DNA position 13170 within *mt-ND5*.

### RNA Methylation Patterns Across Tissues

Across the 39 different tissue/cell types examined, blood, brain, muscle and nerve tissues show the highest levels of RNA methylation across all tRNA P9 sites combined, with average levels of 11-25%, 7-12%, 11%, and 10% respectively (ranges shown where multiple dataset-tissue type pairs are available). In contrast, the lowest levels of tRNA methylation are observed in cell lines, with average levels across P9 sites ranging between 0.3-0.5% in LCLs and at 0.7% in transformed fibroblasts (Fig. 1a, Supplementary Fig. 1). RNA methylation levels also vary between individual tRNA P9 positions along the mitochondrial genome. For example, across all datasets methylation levels at tRNA position 3238 have an average value of 0.9%, whereas the average methylation levels at position 8303 is 12%. To test whether mt-tRNA P9 methylation levels are similar between different P9 sites along the mitochondrial transcriptome within an individual, we measured correlations between methylation levels at each pair of mt-tRNA P9 sites within each dataset-tissue type pair. Across all individuals and dataset-tissue types, all correlation coefficients were significant after correction for multiple testing, and centred around 0.75, ranging between 0.18-0.95 (Fig. 1b), suggesting that methylation levels at different P9 sites are broadly consistent along the mitochondrial transcriptome within each individual.

**Figure 1.**
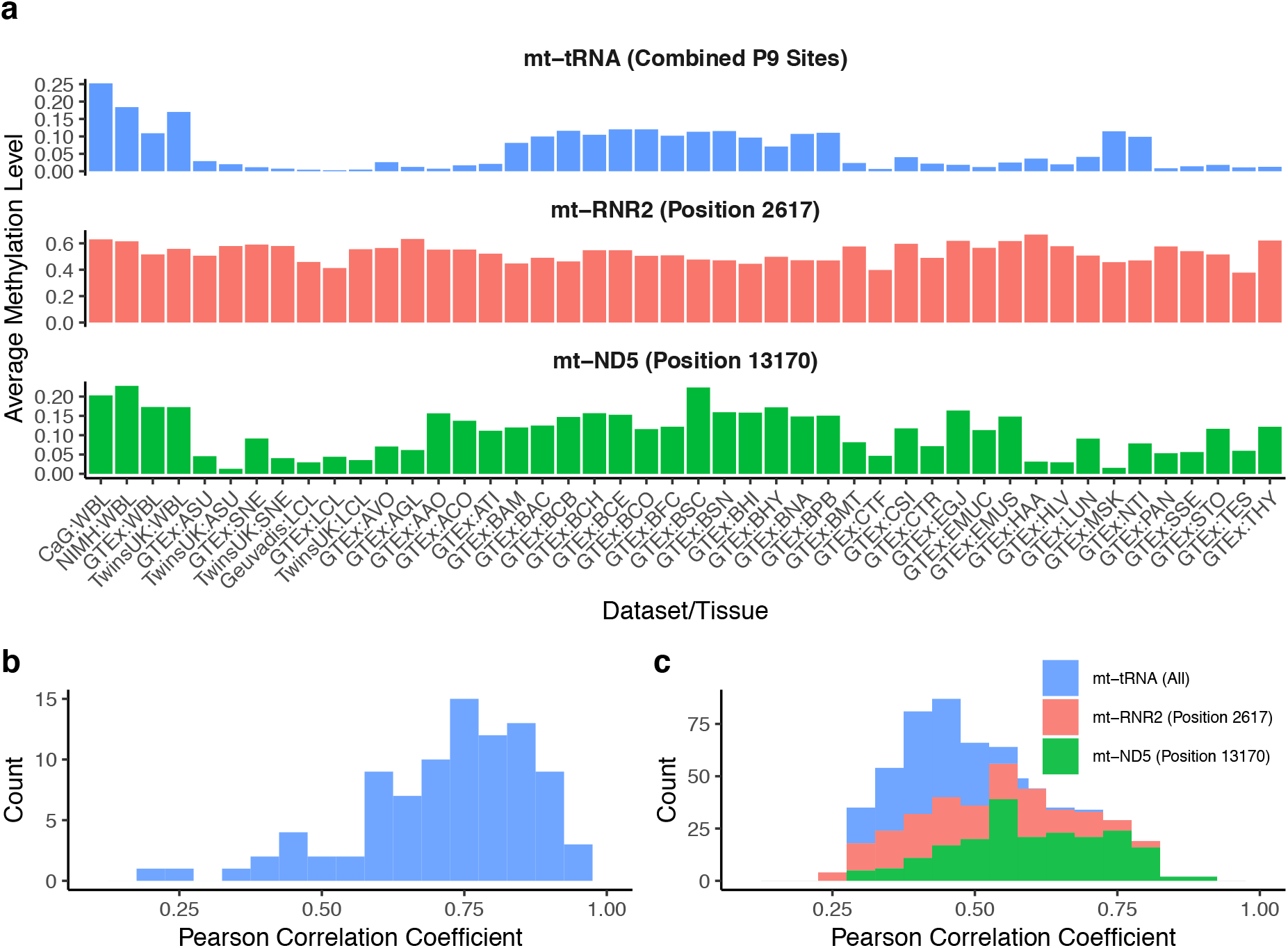
(a) Average RNA methylation levels calculated across datasets and tissue/cell types at three categories of methylated site: averaged values across 11 mt-tRNA P9 sites, mt-RNR2 (position 2617) and mt-ND5 (position 13170). (b) Correlation coefficients between tRNA P9 methylation levels within an individual, measured across individuals in all datasets and tissue types. (c) Correlation coefficients between samples with methylation levels measured in multiple tissues, measured at tRNA P9 sites, mt-RNR2 and mt-ND5. WBL:Whole Blood, ASU:Adipose Subcutaneous, SNE:Skin Not sun Exposed, LCL:Lymphoblastoid Cell Line, AVO:Adipose Visceral Omentum, AGL:Adrenal Gland, AAO:Artery Aorta, ACO:Artery Coronary, ATI:Artery Tibial, BAM:Brain Amygdala, BAC:Brain Anterior cingulate cortex, BCB:Brain Caudate basal ganglia, BCH:Brain Cerebellar Hemisphere, BCE:Brain Cerebellum, BCO:Brain Cortex, BFC:Brain Frontal Cortex, BSC:Brain Spinal cord cervical, BSN:Brain Substantia nigra, BHI:Brain Hippocampus, BHY:Brain Hypothalamus, BNA:Brain Nucleus accumbens basal ganglia, BPB:Brain Putamen basal ganglia, BMT:Breast Mammary Tissue, CTF:Cells Transformed Fibroblasts, CSI:Colon Sigmoid, CTR:Colon Transverse, EGJ:Esophagus Gastroesophageal Junction, EMUC:Esophagus Mucosa, EMUS:Esophagus Muscularis, HAA:Heart Atrial Appendage, HLV:Heart Left Ventricle, LUN:Lung, MSK:Muscle Skeletal, NTI:Nerve Tibial, PAN:Pancreas, SSE:Skin Sun Exposed, STO:Stomach, TES:Testis, THY:Thyroid

Outside of tRNAs, the average methylation level at *mt-rRNR2* (position 2617) and *mt-ND5* (position 13710) transcripts are generally high across all tissues examined (Fig. 1a), with sample wide average values of 55% and 10% respectively. Considering *mt-RNR2* transcript methylation levels across tissues, average levels range between 38% in GTEx Testis and 67% in GTEx Heart (Atrial Appendage). Across tissues, average levels of transcript methylation at *mt-ND5*, a protein coding gene, show more variation, ranging between 1% in TwinsUK subcutaneous adipose and 23% in NIMH whole blood. Within dataset-tissue pairs, we additionally see considerable variation in transcript methylation levels across individuals (Supplementary Fig. 2). For example, at position 2617 in the CARTaGENE whole blood dataset, methylation levels vary between 0.48-0.72, and between 0.11-0.76 in the Geuvadis lymphoblastoid cell line (LCL) data.

To test if mitochondrial RNA methylation levels (at tRNA P9 sites, within the mt-rRNR2 site and within the mt-ND5 site) are correlated across tissue types within an individual, we selected individuals from the GTEx dataset, where RNAseq data from the largest number of alternative tissue types were available, and measured pairwise correlations. For tRNA P9 sites, 11% of pairwise comparisons were significant after correction for multiple testing (median r=0.42, range 0.28-0.7, Fig. 1c). At the *mt-RNR2* (position 2617) and mt-ND5 (position 13710) sites, 76% (median r=0.49, range 0.24-0.81) and 95% (median r=0.59, range 0.3-0.88) of correlation coefficients are significant after correction for multiple testing respectively (Fig. 1c). Collectively, these results demonstrate detectable variation in mitochondrial encoded RNA methylation levels at the individual and population level, as well as consistency in the levels observed along the mitochondrial transcriptome and across tissues, suggesting the presence of shared underlying regulatory mechanisms.

### Nuclear Genetic Associations

To identify nuclear genetic variation associated with mitochondrial-encoded RNA methylation levels, we obtained genome-wide genotyping data for the same samples that we had RNA data for, and carried out association analyses within each of the 39 tissue/cell types for the level of methylation at functionally important positions within the mitochondrial genome. This included methylation levels at: 1a) 13 different tRNA P9 sites along the mitochondrial genome, 1b) a combined measure across multiple tRNA P9 sites (see methods), 2) position 2617 within *mt-rRNR2* and 3) at position 13710 within *mt-ND5*. For tissues where we had multiple independent datasets, which includes whole blood, adipose, skin (non-sun exposed) and LCLs, we then carried out meta-analyses. Across all association studies, we corrected for multiple testing by accounting for genome-wide testing, the number of methylation sites examined and the number of tissues included in the analysis, resulting in a significance threshold of P < 6.79 × 10^−11^.

Across all tissue types and mitochondrial RNA positions where we quantify methylation levels, we find a total of 47 significant associations (peak nuclear genetic variant and mitochondria encoded RNA methylation level pairs). Most associations occur in tissues for which we have multiple independent datasets, and thus larger sample sizes (Table 1, Figure 2 for associations observed in whole blood); 25 nuclear genetic loci are significantly associated with mitochondrial-encoded RNA methylation levels in whole blood, 4 are detected in subcutaneous adipose, 2 in non-sun exposed skin and 1 in LCLs. In single dataset tissues, we identify 15 significantly associated loci across four different tissue types: artery (aorta), nerve (tibial), oesophagus (muscularis and gastroesophageal junction) and heart (left ventricle) (Supplementary Table 2). Across all associated loci, many regions are overlapping in different tissue types and methylation positions; removing all regions that overlap leaves a total of 5 unique regions on the nuclear genome associated with methylation levels at mitochondrial tRNA P9 sites, 2 regions associated with methylation levels at mt-RNR2, 1 region associated with methylation level at both tRNA P9 sites and the mt-RNR2 site and 1 region associated with mt-ND5. Within this, 5 loci represent novel associations detected in this study, and 4 represent replication of previously identified associations in whole blood (Hodgkinson et al. 2014) that we now identify in multiple tissue types across the body.

**Table 1:**
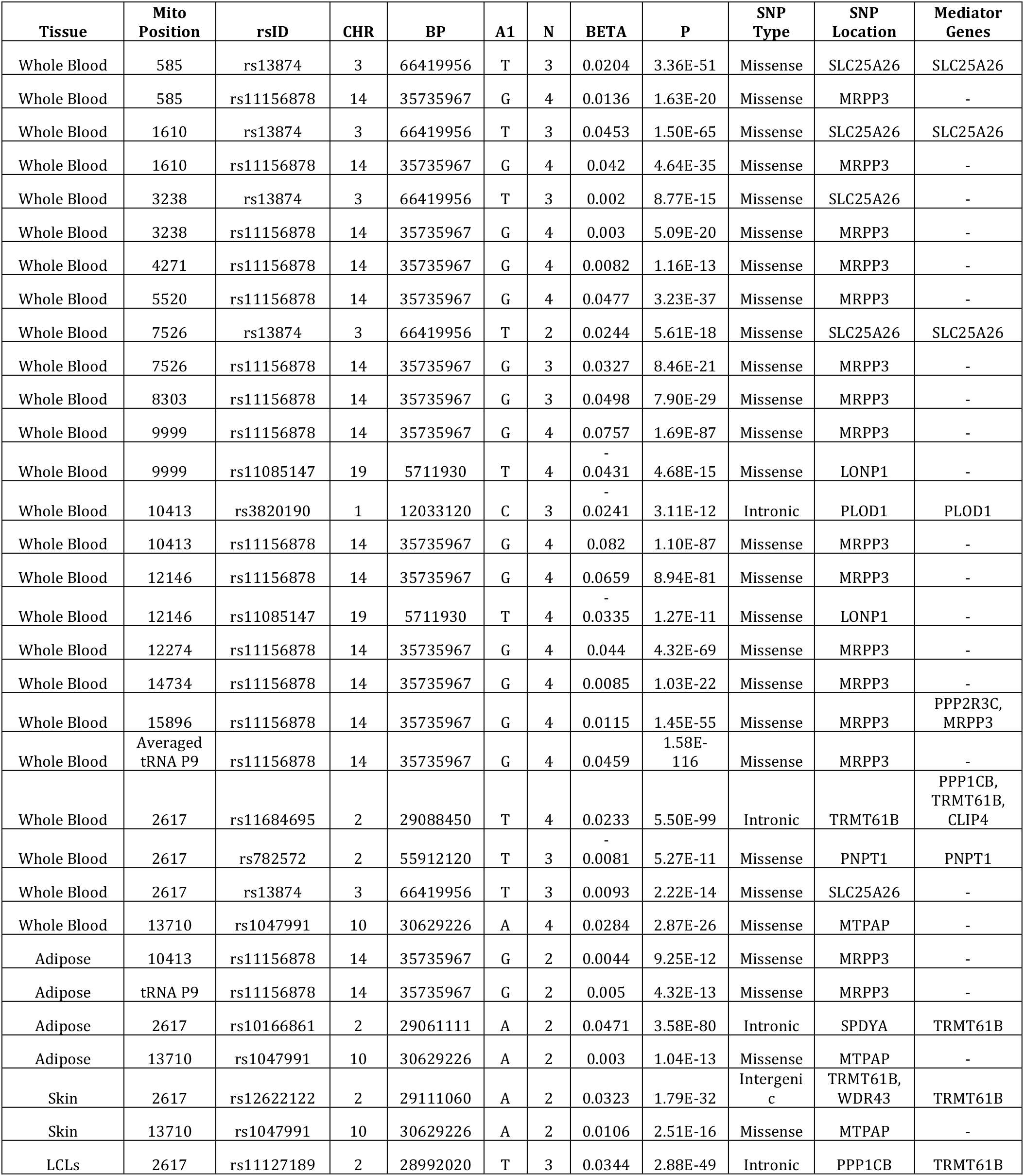
Significant associations between nuclear genetic variation and mitochondrial RNA methylation level for meta-analysed tissues. Alleles presented in the ‘A1’ column represent the minor allele, and the values in column ‘N’ represent the number of studies contributing the meta-analysis, for that row.

**Figure 2.**
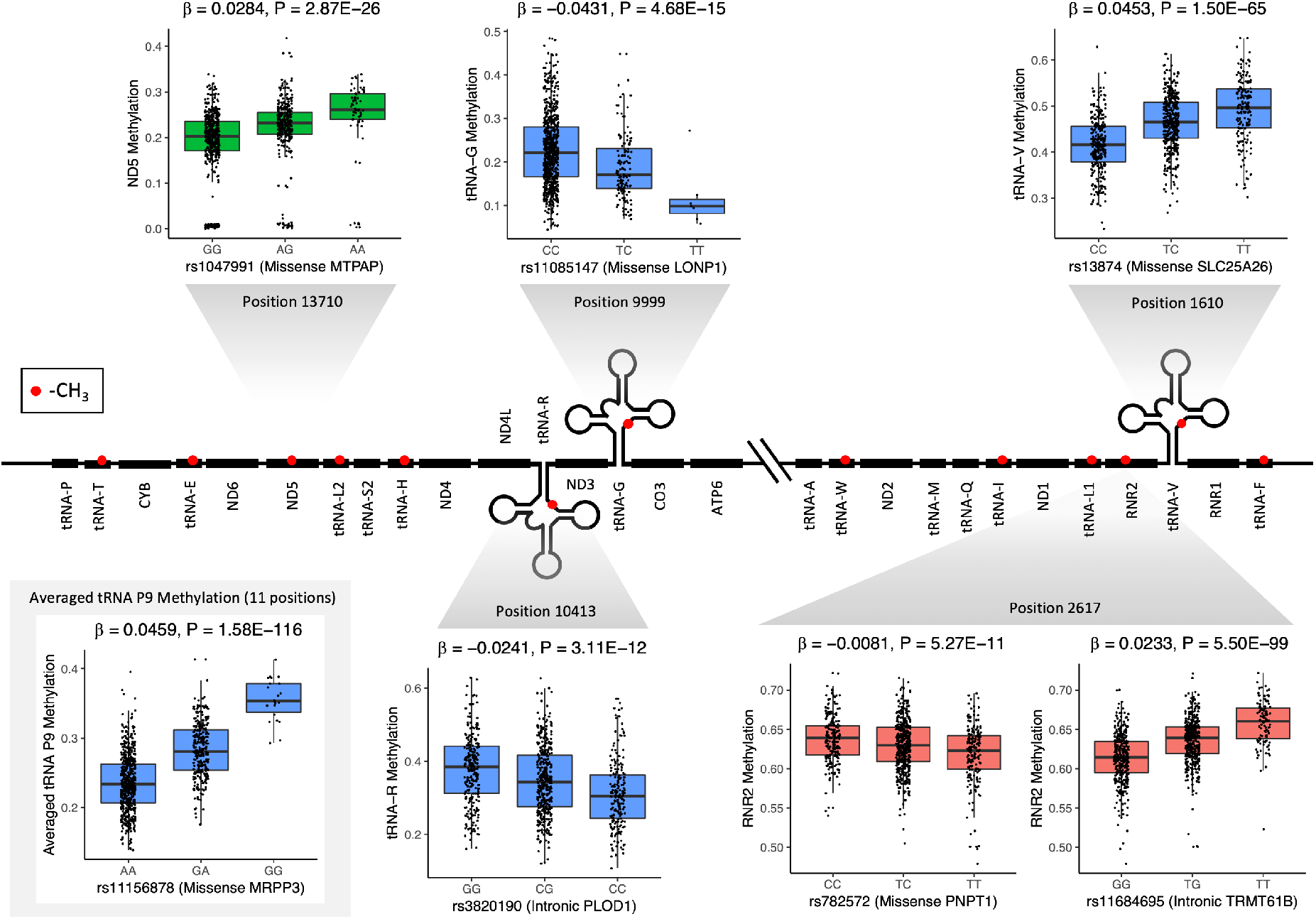
Relationship between genotype and methylation level at multiple positions on the nuclear genome and mitochondrial transcriptome respectively. Methyl groups are represented by red circles along the mitochondrial transcriptome, and methylation levels are shown at three categories of methylated site: at tRNA P9 sites (blue), at position 2617 within *mt-RNR2* transcripts (red) and at position 13170 within *mt-ND5* transcripts (green). Averaged levels of methylation across 11 mt-tRNA P9 sites are additionally shown in the grey shaded box (bottom left). Beta estimates and P-values displayed are from meta-analysis of four independent whole blood datasets, and methylation levels and genotypes displayed in boxplots originate from the CARTaGENE dataset.

### Functional Characterisation

In order to identify the potential genes and mechanisms through which nuclear genetic variants are associated with mitochondrial encoded RNA methylation levels, we prioritised significantly associated variants as potentially causal by annotating them as; 1) resulting in a missense mutation in a protein, 2) affecting expression of a nearby gene and additionally showing evidence of having a mediating effect on mitochondrial RNA methylation level (see Methods), or 3) having the lowest P-value within each associated region.

Applying this approach, we identify a number of novel candidate causal nuclear genetic variants and genes associated with mitochondrial tRNA methylation levels. First, we find a missense mutation (rs11085147) in *LONP1* that is associated with methylation levels at mt-tRNA-G (position 9999) and mt-tRNA-H (position 12146). LONP1 a mitochondrial matrix protein and is involved in degradation of damaged or unfolded polypeptides, in addition to the maturation of certain mitochondrial proteins(Zurita Rendón and Shoubridge 2018). Second, an intronic SNP (rs3820190) in *PLOD1* shows evidence of association with mitochondrial RNA methylation levels at mt-tRNA-R (position 10413) through altered expression of the gene (Table 1). *PLOD1* is not currently thought to be involved in mitochondrial function, however *MFN2* (a gene involved in mitochondrial fusion) was also identified as an eQTL gene for rs3820190. Athough it was not significant in mediation analysis, it remains a viable candidate for the causal gene in this case due to its involvement in mitochondrial processes. Third, within heart tissue (left ventricle), we additionally identify an association between rs10513664, an intronic SNP in the *MECOM* gene, (which has not previously been linked to mitochondria) and methylation level at mt-tRNA-G (position 9999, Supplementary Table 2). MECOM is a transcription factor involved in the proliferation and maintenance of haematopoietic stem cells, and its dysregulated expression has been linked to worsened prognoses in patients with haematological cancers (Shimabe et al. 2009).

Aside from these novel associations, we also replicate previously identified links between missense mutations rs13874 and rs11156878 (within *SLC25A26* and *MRPP3* respectively) and mitochondrial-encoded tRNA methylation levels in whole blood (Hodgkinson et al. 2014). However, we additionally find significant associations between these variants and tRNA P9 methylation levels in adipose, skin, LCLs, artery, oesophagus and nerve tissues (Table 1), showing that these links may be important across the body. *SLC25A26* is a mitochondrial carrier protein, involved in transporting S-adenosylmethionine (a substrate involved with RNA methylation) into mitochondria, and *MRPP3* is part of a complex responsible for 5’ mt-tRNA processing, and thus are directly involved in processes that may impact RNA methylation.

Outside of mitochondrial tRNAs, we identify a novel association between a missense mutation (rs782572) in *PNPT1*, and methylation level at position 2617 within *mt-RNR2*, in addition to an intronic variant (rs11684695) that is significantly associated with transcript methylation levels in whole blood, and mediates mitochondrial RNA methylation levels through the expression of genes including *TRMT61B*, a mitochondrial methyltransferase. At position 13710 within *mt-ND5*, we replicate the association with rs1047991 that was identified in a previous study (Hodgkinson et al. 2014), which is a missense mutation in *MTPAP*, a mitochondrial poly-A polymerase. The latter two associations did not replicate in previous work (Hodgkinson et al. 2014), however do replicate across multiple tissue types in this study (Table 1, Supplementary Table 3), and therefore suggest that these genes may play a global role in the regulation of RNA modification.

### Cross-Tissue Analysis

As whole blood made up our largest dataset (2,424 RNAseq libraries across four datasets), we tested whether nuclear genetic variants associated with mitochondrial RNA methylation levels in whole blood operate in a tissue specific or tissue-wide manner. For each of the 25 significant mitochondrial methylation site-variant pairs in whole blood, we tested for evidence of replication, with the same direction of effect, at matched methylation site and variant pairs in all other tissues. We correct for the number of methylation site-variant pairs we test, and in total 12/25 position-SNP pair associations replicate in at least one other tissue type (Supplementary Table 3). rs11156878, which is associated with the average level of RNA methylation at tRNA P9 sites, replicates in 22 additional tissues. rs11684695, associated with RNA methylation level of position 2617, showed replication in 25 other tissues, and rs1047991, associated with RNA methylation at position 13170, is replicated in 15 additional tissues (Supplementary Table 3). This suggests that certain genetic loci associated with mitochondrial RNA methylation levels in whole blood are active in multiple other tissues, and potentially in a system-wide manner. Other variants however, such as rs13874, which associated with methylation level at multiple individual mt-tRNA P9 sites in whole blood, does not show evidence of association in other tissues, suggesting that it may be tissue specific.

### Consequences of Variation in Mitochondrial RNA Methylation

Methylation modifications at mt-tRNA P9 sites are thought to stabilise the secondary structure of the corresponding mt-tRNA sequences within the mitochondrial transcriptome (Helm et al. 1999), and as tRNA structure may be important for post-transcriptional substrate recognition and cleavage, we tested if methylation levels at tRNA P9 sites were related to mt-mRNA expression. Since RNA methylation levels at P9 sites are strongly associated with a missense mutation in *MRPP3*, which is part of an enzyme (RNase P) that cleaves the 5’ end of mitochondrial tRNAs, we specifically considered the 9 occurrences where an mRNA or rRNA gene is found immediately upstream of a tRNA. Using data from the CARTaGENE project, which is our largest, long-read, paired end dataset, we find that tRNA methylation levels are significantly associated with the expression levels of genes immediately upstream in 6 out of 9 cases at P<0.05 (5 of which show positive correlations) and 3/6 remain significant after correcting for the number of pairs tested (Table 2). This suggests that increased methylation levels at these tRNA P9 sites are associated with increased expression of its nearest upstream gene. To test for replication, we used the GTEx dataset, which is our second largest, long-read, paired end dataset of unrelated samples. We find that 2/6 nominally significant correlations show evidence of replication, with the same direction of effect. These include the positive association between methylation level at *mt-tRNA-L1* and expression of *mt-RNR2*, and the negative relationship between methylation level at *mt-tRNA-G* and expression of *mt-CO3* (Table 2).

**Table 2:**
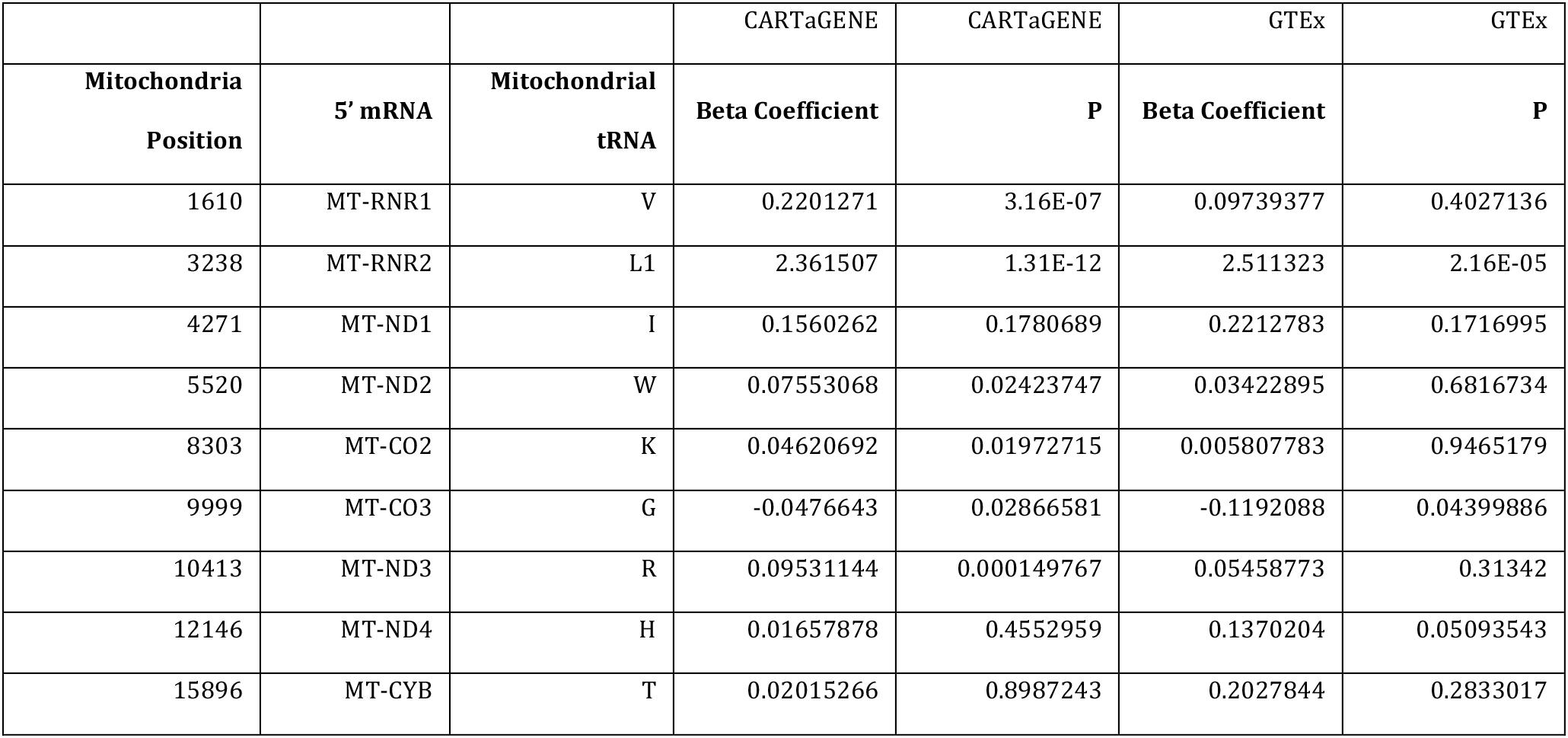
Regression analyses between methylation level at mitochondrial tRNA P9 sites and expression of its 5’ mRNA, in whole blood datasets.

### Overlap with Disease Associated Loci

Finally, to identify potential links between genetic variants associated with mitochondrial RNA methylation levels and complex-traits and phenotypes, we looked for overlaps between peak associated variants (and SNPs in strong LD, r^2^ ≥ 0.8), and genome-wide significant disease associated variants in the NHGRI-EBI GWAS Catalog (MacArthur et al. 2017). We find overlaps with diastolic (Ehret et al. 2016) and systolic blood pressure (Kristiansson et al. 2012; Ehret et al. 2016), Moyamoya disease (Duan et al. 2018), estrogen-receptor negative breast cancer (Couch et al. 2016; Milne et al. 2017) and adolescent idiopathic scoliosis (Liu et al. 2018) (Supplementary Table 4).

First, the peak genetic variant (rs3820190) associated with methylation at *mt-tRNA-R* (position 10413) through the expression of *PLOD1*, is in high LD with a genetic variant (rs2273291) linked to Moyamoya disease, a rare cerebrovascular disease that is characterised by progressive narrowing of the lower brain vasculature that can lead to transient ischemic attacks and stroke (Duan et al. 2018). Duan *et al* (2018) also report that a subset of individuals with Moyamoya disease (that also carry the risk allele at rs2273291) have slightly elevated levels of serum homocysteine, which itself is associated with an increased risk of stroke (Wald et al. 2002). Interestingly, S-adenosylmethionine is an intermediary substrate in the homocysteine biosynthesis pathway (Hankey et al. 2004), possibly explaining the link observed in the present study, as an increased level of homocysteine could mean that there is less S-adenosylmethionine available for RNA methylation.

Second, the missense mutation in *PNPT1* (rs782572) that is associated with methylation level of mt-RNR2 (position 2617), is in high LD with rs197548, which is associated with systolic and diastolic blood pressure (Ehret et al. 2016). Mitochondria have previously been linked to increased blood pressure, predominantly through mechanisms involving mitochondrial oxidative stress, however the exact mechanisms by which this is the case remain unclear (Dikalov and Dikalova 2016). We have previously observed associations between nuclear genetic variants influencing the expression of certain mitochondrial genes, mediated through the expression of *PNPT1* (Ali et al. 2019), and the identification of the overlap here suggests that RNA methylation level may also be playing a contributory role in the process. A SNP within an intron of the *MECOM* gene (rs6779380) has previously been associated with changes in blood pressure (Ehret et al. 2016), however this variant it is not in LD with the variant that we identify in heart (left ventricle, rs10513664), despite the latter also being located within an intron of *MECOM*.

Finally, rs11684695, which is associated with methylation level at mt-*RNR2* (position 2617), through the expression of *TRMT61B*, is in high LD with rs4577244 and rs67073037, both of which have been linked to breast cancer. The role of mitochondria in cancer has been debated since the discovery of the Warburg effect (WARBURG 1956), and subsequent research has linked many additional pathways/features of the mitochondria to tumorigenesis, including through alterations in its roles in cell death, metabolism, and oxidative stress (Vyas et al. 2016). In previous work, we observed an increase in the level of tRNA P9 methylation levels in cancer tumours vs matched normal tissues (Idaghdour and Hodgkinson 2017); our observation of an overlap here suggests that methylation level at mt-*RNR2* may also be involved. Additionally, rs11684695 is also in high LD with rs6737027, which has been associated with adolescent idiopathic scoliosis (Liu et al. 2018), and an increased incidence of scoliosis in patients with mitochondrial myopathies has also been reported (Li et al. 2015).

## Discussion

RNA modifications represent an additional layer of control in the regulation of gene expression. They are found extensively throughout both the nuclear and mitochondrial transcriptome, where they play important roles in structural stability and translation efficiency. Using mitochondria as a model system, we characterise RNA methylation levels at multiple functionally important sites on the mitochondrial transcriptome, across a total of 39 tissues/cell types. We find that RNA methylation levels are correlated along the transcriptome, but vary between tissues, with blood and brain tissues showing the highest levels of variation. As the mitochondrial and nuclear genomes have co-evolved over evolutionary time, we also link variation in mitochondrial RNA methylation levels to genetic variation in the nuclear genome.

In total, we associate 8 nuclear genes to fundamental biological processes taking place in human mitochondria. Within this, we identify novel associations between mitochondrial RNA methylation levels and missense mutations in *LONP1* and *PNPT1*, eQTLs regulating the expression of *PLOD1* and *TRMT61B* and an intronic variant within *MECOM*. Furthermore, we find that previously identified associations (Hodgkinson et al. 2014), which have been linked to *MRPP3*, *TRMT61B* and *MTPAP* (rs11156878, rs11684695 and rs1047991 respectively), occur in multiple tissue types, which may have important implications for disease.

*MRPP3* is the catalytic subunit of mitochondrial RNase P, a complex responsible for the 5’ cleavage of mt-tRNAs (Holzmann et al. 2008), and is active only in the presence of its other subunits (Reinhard et al. 2015), MRPP1 and MRPP2. The MRPP1 and MRPP2 sub-complex however, is able to carry out its methyltransferase activity independently of MRPP3 (Vilardo et al. 2012), so the association between rs11156878 within *MRPP3* with methylation of mt-tRNAs is likely detected due to its effect on cleavage capacity. *TRMT61B* is a methyl-transferase that is responsible for the methylation of position 58 on certain mitochondrial tRNAs, in addition to position 2617 in *mt-RNR2* (Bar-Yaacov et al. 2016). In the present study, an intronic SNP (rs11684695) is associated with increased levels of methylation at *mt-RNR2*, as well as increased expression of *TRMT61B*, likely explaining the relationship detected here. *MTPAP* is responsible for the polyadenylation of the 3’ end of mitochondrial mRNAs, and this polyadenylation is important for the stability of the transcripts (Nagaike et al. 2005), which potentially influences the methylation levels of *mt-ND5* transcripts. Of genes newly implicated with mitochondrial RNA methylation levels through missense mutations, *LONP1* has been shown to degrade *MRPP3* as part of the mitochondrial unfolded protein response (Münch and Harper 2016), possibly explaining its association with methylation level. Finally, *PNPT1* is involved in multiple metabolic RNA processes in mitochondria, and reduction of *PNPT1* levels results in impaired mitochondrial processing and accumulation of large polycistronic transcripts, possibly due to its connection to the import of RNase P RNA into the mitochondria (Wang et al. 2010a; von Ameln et al. 2012), again likely explaining why it is associated with methylation level in this study.

Interestingly, rs13874, a missense mutation in *SLC25A26*, tends to only be associated with tRNAs towards the beginning of the mitochondrial transcript in whole blood, and does not show evidence of replicating across tissues. *SLC25A26* is a mitochondrial carrier protein that is responsible for transporting S-adenosylmethionine into the mitochondria, which is a methyl group donor in methylation reactions. The absence of rs13874 replication across tissues may be related to the fact that the highest levels of methylation are seen in whole blood, in combination with *SLC25A26* concurrently having the lowest levels of expression in whole blood (https://gtexportal.org/home/gene/SLC25A26). Therefore, the effect of a possible transportation deficiency may only be observed in tissues where the requirement for methylation is high. As blood is the tissue in which we detect the highest levels of methylation, but also the lowest levels of *SLC25A26* expression, this mutation may have particularly important implications for blood-based processes and diseases.

The downstream functional consequences of altered mitochondrial RNA processing are well documented in human cell lines and model organisms (Sanchez et al. 2011; Perks et al. 2017), but here using *in vivo* data we show that natural variation of mitochondrial RNA methylation levels in ‘healthy’ individuals may influence mitochondrial processes (namely changes in mitochondrial gene expression). Disruption or perturbation of the function of nuclear genes that we have implicated in mitochondrial RNA methylation can have serious phenotypic consequences. In humans for example, a mutation in *PNPT1* has also been linked with impaired import of RNA into the mitochondria, and leads to Combined Oxidative Phosphorylation Deficiency (Vedrenne et al. 2012) and also Autosomal Recessive Deafness (von Ameln et al. 2012); missense mutations in *LONP1* have also been implicated with CODAS syndrome, which is a developmental disorder affecting multiple systems (cerebral, ocular, dental, auricular and skeletal) (Strauss et al. 2015); mutations in *SLC25A26* have been linked to Combined Oxidative Phosphorylation Deficiency (Kishita et al. 2015) and mutations in *MTPAP* have been linked to spastic ataxia (Crosby et al. 2010). Similarly, *TRMT61B* has been identified as a differentially expressed gene in a small cohort of Alzheimer’s disease cases, when compared to matched controls (Perks et al. 2017), and knockdown of the drosophila homolog of *MRPP3* leads to the loss of locomotive function in *Drosophila*, similarly to what is seen in Parkinson’s disease (Sen et al. 2016).

These examples are phenotypically varied and are the result of extreme alterations in the function of the corresponding gene. The genetic variation associated with RNA methylation levels in this study have less extreme effects, however, they are linked with changes in post-transcriptional processing and downstream expression of mitochondrial genes. While this variation might be tolerated under normal physiological situations, the introduction of stressful situations, for example during increased mitochondrial damage through ageing, may be enough to push a tissue that is heavily reliant on appropriate mitochondrial function into dysfunction. Conversely, altered mitochondrial expression over a long duration may be enough to lead to negative later life consequences. Impaired mitochondrial gene expression due to the heterozygous knockout of *PTCD1*, a mitochondrial RNA processing enzyme for example, has been linked to later-life obesity in mice (Perks et al. 2017). In this study, we find overlaps between genetic variants associated with mitochondrial RNA methylation levels and variants linked to blood pressure, breast cancer and Moyamoya disease, suggesting that altered mitochondrial RNA modification may play a role in more complex diseases. Overall, as the expression of mtDNA is regulated primarily at the post-transcriptional level, it is important to understand how variation in nuclear encoded genes affects mitochondrial RNA methylation and gene expression, as changes in this can lead to changes in mitochondrial metabolism that impact mitochondrial function.

## Methods

### Data Description

RNA sequence and genotype data were obtained from five independent, publically available projects, including:

CARTaGENE (Awadalla et al. 2013): A population-based cohort comprised of people aged between 40-69, from Quebec, Canada. Whole blood samples were taken for RNA sequencing and genotyping, producing 100bp paired-end RNAseq reads and genotypes from the Illumina Omni2.5M genotyping array. Samples with RNAseq data from multiple sequencing runs, that passed quality control, were merged before the alignment stage. Data were obtained through application to the data access committee (instructions are available at www.cartagene.qc.ca).

NIMH (National Institute of Mental Health) Genomics Resource (Battle et al. 2014): Whole blood samples were collected for RNA sequencing and genotyping from the Depression Genes and Networks study. Individuals were aged 21-60, and are a case/control cohort. 50bp single-end RNAseq reads were produced, along with genotypes from the Illumina HumanOmni1-Quad BeadChip. Mapped RNAseq reads for duplicate samples that passed quality control were merged for further analysis, and samples failing QC were discarded. Data were obtained after application to the data access committee (through www.nimhgenetics.org)

Geuvdais Project (Lappalainen et al. 2013): LCL samples from the 1000 Genomes cohort were RNA sequenced to produce 75bp paired-end RNAseq reads and were obtained from the European Nucleotide Archive under submission number ERA169774. Mapped DNA sequence data from phase 1 of the project were downloaded from the 1000 Genomes FTP site (v5a.20130502).

TwinsUK Project (Grundberg et al. 2012): Female monozygotic twin pairs, dizygotic twin pairs and singletons, aged between 38-85 were recruited for RNA sequencing and genotyping. Biopsies from subcutaneous adipose tissue and skin were collected, as well as peripheral blood samples for additional generation of lymphoblastoid cell lines (LCLs). 50bp paired-end RNAseq data were produced from these tissues as well as genotypes from Illumina HumanHap300 and Illumina HumanHap610Q genotyping arrays. Data were obtained through application to the TwinsUK data access committee and then downloaded from the European Genome-Phenome archive (https://ega-archive.org) through study ID EGAS00001000805.

GTEx (Genotype-Tissue Expression) Project (Consortium 2013): Multiple tissue samples were collected from deceased individuals for RNA sequence analysis and dense genotyping, with sample age range varying between 20-71. We use a combination of data from the pilot and midpoint phases of the GTEx project, where samples were genotyped in the Illumina Omni5M and Illumina Omni2.5M genotyping arrays respectively. RNAseq read lengths produced by the project varied, and we analyse samples with 75bp long reads only. Data were obtained by application to dbGaP through accession number phs000424.v6.p1.

Full data accession information, sample sizes and tissue types are described further in Supplementary Table 1.

### RNAseq Mapping

FastQC [v0.11.3] (https://www.bioinformatics.babraham.ac.uk/projects/fastqc/) was run on raw RNAseq data, and samples with drops in base quality below phred 20 or uncalled bases in the middle of reads were discarded. RNAseq reads were then pre-processed to remove adaptor sequences and low quality trailing bases (Phred < 20) using TrimGalore [v0.4.0] (https://www.bioinformatics.babraham.ac.uk/projects/trim_galore/). Poly-A/T sequences greater than 4bp were also removed from read termini using PRINSEQ-lite [v 0.20.4] (Schmieder and Edwards 2011). Remaining reads with >20 nucleotides were then mapped to the human reference sequence (1000G GRCh37 reference, which contains the mitochondrial rCRS NC_012920.1) using STAR [2.5.2a_modified] 2-Pass mapping, allowing approximately 1 mismatch per 18 bases per read, rounded down to the nearest integer. STAR soft-clipping was also allowed. After mapping, FastQC [v0.11.3] was rerun on data, and samples with median sequence quality scores falling below Phred 20 were removed from further analysis. SAMtools [v1.4.1] (Li et al. 2009) was then used to retain only properly paired and uniquely mapped reads. This stringent step was applied to ensure that analysed reads originated from the mitochondria, rather than nuclear encoded fragments of mitochondrial DNA (NUMTs). Transcript abundances were calculated using the ‘intersection non-empty model’ within HTseq [v0.6.0] (Anders et al. 2015), and gene expression counts were then quantified according to transcripts per million (TPM). Genes expressed in all samples with an average TPM value > 2 were used to calculate principal components (PCs) in R and outliers identified from visualisation of PC1, PC2 and PC3 were excluded. Samples were further excluded for having: fewer than 5,000,000 remaining reads, fewer than 10,000 mitochondrial reads, rRNA content greater than 30%, RNAseq mismatch percentage greater than 1%, or intergenic read percentage greater than 30%.

### Quality Control, Phasing and Imputation of Genotype Data

QTLtools [v1.0] (https://qtltools.github.io/qtltools/) was used to ensure sample labelling was consistent between genotype and RNAseq data. Quality control (QC) of genotype data was carried out using PLINK [v1.90b3.44] (Chang et al. 2015). Duplicate samples, genetic PC outliers, samples with unexpected relatedness and samples with outlying heterozygosity rates were removed, in addition to samples with discrepant reported and genotypic sex information, or ambiguous X chromosome homozygosity estimates. Samples with > 5% missing genotype data were also excluded. SNPs were removed for violating Hardy-Weinberg Equilibrium (HWE) with a P-value < 0.001, for having a genotype missingness > 5% or for having a minor allele frequency (MAF) < 1%. SNPs coded according to the negative strand were flipped to the positive strand. SNPs remaining on autosomal chromosomes were phased using default settings within SHAPEIT [v2.r837] (Delaneau et al. 2013), for all datasets with array genotype data. Phased chromosomes were imputed in 2Mb intervals using default settings within IMPUTE2 [v2.3.2] using 1000 Genomes Phase 3 individuals as a reference population (Marchini and Howie 2010; Howie et al. 2011). Imputed genotypes were then hard called with a minimum calling threshold of 0.9 using GTOOL [v0.7.5] (http://www.well.ox.ac.uk/~cfreeman/software/gwas/gtool.html) and filtered out for having an IMPUTE2 info score < 0.8, MAF < 5%, genotype missingness > 5%, HWE P < 0.001 or for being multi-allelic. Datasets genotyped on two different arrays were imputed separately and then merged.

### Quantification of Mismatch Rate at Modified Sites

Previous studies have shown that the proportion of mismatching bases at certain sites on the mitochondrial transcriptome can be used to represent the level of post-transcriptional methylation at these sites. During library preparation for RNA sequencing, methylation modifications on transcripts can interfere with the reverse transcription process by causing the reverse transcriptase to randomly incorporate nucleotide bases at the methylated position (Hauenschild et al. 2015). Though not a direct reflection of methylation level (Hauenschild et al. 2015), this mismatch signature can be used to estimate the level of methylation present on transcripts by measuring the proportion of non-reference alleles at modified sites (Mercer et al. 2011; Sanchez et al. 2011; Hodgkinson et al. 2014; Idaghdour and Hodgkinson 2017). Previous work has demonstrated that methylation levels estimated this way are repeatable across experiments (Hodgkinson et al. 2014), and comparison of samples treated with demethylation enzymes to untreated controls confirms the presence of methylation at the ninth position of 19/22 mt-tRNA positions, at similar levels to when measured by primer extension (Clark et al. 2016). Here, we consistently detect modification levels at levels ≥ 1% at 13/19 of these mt-tRNA sites, which correspond to the following mitochondrial genomic coordinates: 585, 1610, 3238, 4271, 5520, 7526, 8303, 9999, 10413, 12146, 12274, 14734 and 15896. We also calculate methylation level at mtDNA positions 2617 and 13710, which correspond to locations within mt-RNR2 and mt-ND5 respectively; methylation levels at these sites can also be determined using RNAseq (Hauenschild et al. 2015; Li et al. 2017).

Positional read coverage at mitochondrial sites were summarised using SAMtools mpileup and mismatch rate was calculated from sites with a nucleotide quality score ≥ Phred 13 and coverage ≥ 20x. A site was then considered to show evidence of being methylated if the average proportion of mismatches within a dataset was greater than 1%, as below this level, mismatched due to the presence of methylation is indistinguishable from mismatches due to sequencing error. A combined measure of methylation level was also calculated by averaging across 11 mt-tRNA p9 sites: 585, 1610, 4271, 5520, 7526, 8303, 9999, 10413, 12146, 12274 and 14734 (where values are present), in order to gain an idea of processes influencing post-transcriptional methylation overall. These sites consistently show variation in whole blood data and have previously been used as an estimate of combined methylation (Hodgkinson et al. 2014; Idaghdour and Hodgkinson 2017). Methylation values 3 standard deviations from the mean were masked to avoid association results being driven by extreme values.

Pearson’s correlation coefficients between methylation levels at different tRNA P9 sites (including averaged levels across P9 sites) were calculated within individuals, across all datasets and tissue types available. A total of 91 comparisons that were carried out. Pearson’s correlation coefficients between methylation level at the same position, across multiple tissues, were carried out using measurements from the GTEx dataset. For a position to be compared between tissues, we required that both tissues have an average methylation level of 1% and at least 100 pairs of data points to compare. For tRNA P9 sites, there are 1657 comparisons, for positions 2617 and 13710 there are 219 comparisons.

### Association Analyses and Meta-Analysis

Association analyses studies were carried out for modification positions with an average methylation level ≥ 1% variation per dataset. Association analyses were carried out separately for each position and tissue, using linear models in PLINK [v1.9] (Chang et al. 2015). For the TwinsUK tissues, GEMMA [v0.96] (Zhou and Stephens 2012) was used to calculate a relatedness matrices and association tests were carried out using univariate linear mixed models. Covariates used in the association model included 5 study specific genetic PCs and 10 PEER factors calculated from RNAseq data using PEER [v1.0] (Stegle et al. 2012) for tissues with ≥100 samples or 5 genetic PCs and 5 PEER factors for tissues with <100 samples. Additional covariates included in the association model were sex, genotyping array and RNA-sequencing batch information, where available and where relevant. Tissues with multiple datasets were meta-analysed using PLINK [v1.9], under a fixed effects model.

### Cis-eQTL Identification and Mediation Analysis

To identify genes through which nuclear genetic variants associated with mitochondrial post-transcriptional methylation levels were acting, we carried out a cis-eQTL analysis, followed by mediation analysis. Cis-eQTLs and eGenes were identified by selecting protein coding genes within a 1Mb interval of peak SNPs and testing for association between the genotype at the peak SNP and cis-gene expression level in the corresponding tissue type, using quantile normalised RNAseq data. For tissues with multiple datasets, cis-eQTLs were identified in the dataset with the largest sample size. For blood, this corresponded to the CARTaGENE dataset, for adipose and skin tissues, this corresponded to unrelated samples in the TwinsUK dataset, and for LCLs, this corresponded to the Geuvadis dataset. eGENEs are reported if significant in any corresponding tissue type and tests were corrected for the number of cis-genes tested for association, per peak SNP. To identify causal relationships between genotype, gene expression levels and methylation levels, we tested for significant mediation effects using 1000 bootstrapping simulations with the ‘Mediation’ package in R. Mediation analyses were carried out using quantile normalised gene expression values, in the same datasets that eQTLs and gene pairs were identified in. Each mediation test was corrected for the number of significant cis-genes tested for evidence of a mediation effect with a SNP.

### Candidate Causal Variant Selection

Significant SNPs were defined as SNPs passing genome-wide significance after correction for the number of methylation positions analysed, and the number of unique dataset-tissue type pairs they were analysed in, resulting in a threshold of (6.79 × 10^−11^). SNPs passing corrected genome-wide significance were annotated using the ‘geneanno’ option in ANNOVAR [v2017-07-17] (Wang et al. 2010b). Candidate causal SNPs were further shortlisted as corrected genome-wide significant SNPs that resulted in a missense mutation, or showed evidence for mediating methylation levels as a cis-eQTL, or for having the lowest P-value.

### Cross Tissue Replication Analysis

Peak methylation site-variant pairs from the whole blood meta-analysis were tested for nominal evidence of significance in matched methylation site-variant pairs in other tissue types (excluding the tissues that were meta-analysed). Nominal significance is defined as P<0.05, corrected for the number of tissues the SNP-trait pair is tested in, and the number of SNP-trait pairs tested.

### Consequences of Variation in Mitochondrial RNA Methylation Levels

To test if methylation level at mitochondrial tRNA P9 sites have an impact on expression levels of immediately upstream mitochondrial genes, we regressed the expression levels of the upstream gene on the level of methylation at the relevant tRNA P9 site, including batch, gPC 1-5 and PEER factors 1-10 as covariates in the model. This analysis was carried out in the CARTaGENE whole blood dataset, which is our largest, paired-end, long read dataset, and replicated in the GTEx whole blood dataset. We tested 9 regions of the mitochondrial genome, where there is an rRNA or mRNA gene immediately upstream of a tRNA gene and identified significant relationships at P<0.05, corrected for by the number of positions tested.

### Overlap with Disease

We additionally tested for overlap between genetic variants identified in our association studies, and SNPs in strong LD (r^2^ ≥ 0.8) within a 500kb interval, with genome-wide significant SNPs associated with disease phenotypes reported in the NHGRI-EBI GWAS Catalog (MacArthur et al. 2017). Variants in strong LD with peak SNPs were identified from the NIMH, CARTaGENE, TwinsUK (unrelated samples only), GTEx and Geuvadis datasets using PLINK, and were overlapped with data present in the NHGRI-EBI GWAS Catalog in March 2019.

## Acknowledgements

We thank the CARTaGENE platform, GTEx, TwinsUK, the NIMH Genomics Resource and the Geuvadis project for use of RNA sequencing data. A.H. holds a Medical Research Council (MRC) eMedLab Medical Bioinformatics Career Development Fellowship, funded from award MR/L016311/1. A.H. also holds a WHRI-Academy Marie Curie (COFUND) Fellowship and the research leading to these results has received funding from the People Programme (Marie Curie Actions) of the European Union’s Seventh Framework Programme (FP7/2007-2013) under REA grant agreement n° 608765. Work presented here reflects only the author’s views and not the views of the European Commission. A.T.A. is supported by the Generation Trust. Y.I. is funded by a New York University Abu Dhabi research grant (AD105). The research was supported by the National Institute for Health Research (NIHR) Biomedical Research Centre based at Guy’s and St Thomas’ NHS Foundation Trust and King’s College London. The views expressed are those of the authors and not necessarily those of the NHS, the NIHR or the Department of Health. For GTEx data: the Genotype-Tissue Expression (GTEx) Project was supported by the Common Fund of the Office of the Director of the National Institutes of Health (commonfund.nih.gov/GTEx). Additional funds were provided by the NCI, NHGRI, NHLBI, NIDA, NIMH, and NINDS. Donors were enrolled at Biospecimen Source Sites funded by NCI\Leidos Biomedical Research, Inc. subcontracts to the National Disease Research Interchange (10XS170), Roswell Park Cancer Institute (10XS171), and Science Care, Inc. (X10S172). The Laboratory, Data Analysis, and Coordinating Center (LDACC) was funded through a contract (HHSN268201000029C) to the The Broad Institute, Inc. Biorepository operations were funded through a Leidos Biomedical Research, Inc. subcontract to Van Andel Research Institute (10ST1035). Additional data repository and project management were provided by Leidos Biomedical Research, Inc.(HHSN261200800001E). The Brain Bank was supported supplements to University of Miami grant DA006227. Statistical Methods development grants were made to the University of Geneva (MH090941 & MH101814), the University of Chicago (MH090951,MH090937, MH101825, & MH101820), the University of North Carolina - Chapel Hill (MH090936), North Carolina State University (MH101819),Harvard University (MH090948), Stanford University (MH101782), Washington University (MH101810), and to the University of Pennsylvania (MH101822). For NIMH data: data was provided by Dr. Douglas F. Levinson. We gratefully acknowledge the resources were supported by National Institutes of Health/National Institute of Mental Health Grants 5RC2MH089916 (PI: Douglas F. Levinson, M.D.; Co-investigators: Myrna M. Weissman, Ph.D., James B. Potash, M.D., MPH, Daphne Koller, Ph.D., and Alexander E. Urban, Ph.D.) and 3R01MH090941 (Co-investigator: Daphne Koller, Ph.D.). For TwinsUK data: The TwinsUK study was funded by the Wellcome Trust and European Community’s Seventh Framework Programme (FP7/2007-2013). The TwinsUK study also receives support from the National Institute for Health Research (NIHR)-funded BioResource, Clinical Research Facility and Biomedical Research Centre based at Guy’s and St Thomas’ NHS Foundation Trust in partnership with King’s College London.

## Author Contributions

A.H. and Y.I. designed the study. A.A. processed raw data and performed QC and analyses. A.A., Y.I. and A.H. wrote the paper.

